# Triplanar ensemble U-Net model for white matter hyperintensities segmentation on MR images

**DOI:** 10.1101/2020.07.24.219485

**Authors:** Vaanathi Sundaresan, Giovanna Zamboni, Peter M. Rothwell, Mark Jenkinson, Ludovica Griffanti

## Abstract

White matter hyperintensities (WMHs) have been associated with various cerebrovascular and neurodegenerative diseases. Reliable quantification of WMHs is essential for understanding their clinical impact in normal and pathological populations. Automated segmentation of WMHs is highly challenging due to heterogeneity in WMH characteristics between deep and periventricular white matter, presence of artefacts and differences in the pathology and demographics of populations. In this work, we propose an ensemble triplanar network that combines the predictions from three different planes of brain MR images to provide an accurate WMH segmentation. Also, the network uses anatomical information regarding WMH spatial distribution in loss functions for improving the efficiency of segmentation and to overcome the contrast variations between deep and periventricular WMHs. We evaluated our method on 5 datasets, of which 3 are part of a publicly available dataset (training data for MICCAI WMH Segmentation Challenge 2017 - MWSC 2017) consisting of subjects from three different cohorts. On evaluating our method separately in deep and periventricular regions, we observed robust and comparable performance in both regions. Our method performed better than most of the existing methods, including FSL BIANCA, and on par with the top ranking deep learning method of MWSC 2017.

## 1. Introduction

White matter hyperintensities (WMHs) of presumed vascular origin appear as bright localised areas in T2-weighted and fluid-attenuated inversion recovery (FLAIR) images (Wardlaw et al., 2013), and could appear hypointense on T1-weighted images. WMHs occur commonly in patients with cerebrovascular diseases (Li et al., 2013; Simoni et al., 2012) and have been associated with cognitive decline, atrophy and neurodegenerative diseases such as dementia (Debette et al., 2010; Prins and Scheltens, 2015; Pantoni et al., 2005). However, they are also found commonly in healthy elderly subjects (Zamboni et al., 2019). Therefore, the relationship between the occurrence of WMHs and various clinical factors is not yet fully understood. While various visual rating scales are available (Fazekas et al., 1987; Scheltens et al., 1993; Wahlund et al., 2001) and can provide qualitative or categorical information regarding WMHs, they provide limited information regarding the spatial distribution of WMHs. Voxel-wise WMH maps, on the other hand, enable more precise quantification of WMHs and open up the possibility of studying the relationship between the spatial location/distribution of WMHs and various clinical factors. In turn, this helps to identify patterns of normal and pathological ageing (Rostrup et al., 2012; Biesbroek et al., 2013). Hence, location and volume-based lesion characterisation are being increasingly considered in clinical setting (Wardlaw et al., 2013; Smith et al., 2019). Given the importance of analysing the clinical impact of WMHs, especially in large cohorts, manual segmentation of WMHs is time consuming and is prone to intra/inter-rater variability. Hence, an automated method to provide exact voxel-level localisation and accurate quantification of WMHs would be highly useful.

Several automated WMH segmentation methods have been proposed (Caligiuri et al., 2015) using features based on intensity (Ong et al., 2012; Damangir et al., 2012), combined with anatomy (Ong et al., 2012; Damangir et al., 2012) and appearance (such as shape, contrast etc) (Samaille et al., 2012; Griffanti et al., 2016). Among the existing methods using hand-crafted features, unsupervised methods such as clustering (Admiraal-Behloul et al., 2005; Ong et al., 2012; Samaille et al., 2012), supervised classification algorithms (Anbeek et al., 2004; Damangir et al., 2012; Yoo et al., 2014; Griffanti et al., 2016) and probabilistic approaches (Yang et al., 2010) have been proposed. Despite the large amount of methods developed for WMH segmentation, only a few of them are publicly available (Damangir et al., 2012; Lao et al., 2008; Schmidt et al., 2012; Griffanti et al., 2016). Using hand-crafted features might not be sufficient to capture the lesion patterns and to overcome noise and artefacts. The lesion characteristics of WMHs show high variability depending on their location, making their segmentation challenging. For instance, between periventricular WMHs (PWMHs) and deep WMHs (DWMHs) (Griffanti et al., 2017), PWMHs usually appear brighter, larger and often form confluent lesions, with higher contrast when compared to DWMHs that usually occur as small punctate lesions. Also, the lesion load and distribution are influenced by demographic factors (e.g. age) and clinical conditions (e.g. cognitive decline). Additionally, artefacts (e.g. Gibbs ringing, motion artefacts) and noise that occur during image acquisition also affect the segmentation performance.

Deep learning (DL) allows computational models with multiple layers to learn data representations at different layers of abstraction (LeCun et al., 2015), thus utilising more contextual information compared to the hand-crafted features. With increase in computational resources, such as graphical processing units (GPUs) and techniques such as data augmentation (Russakovsky et al., 2015), DL has been quickly emerging as a reliable segmentation tool in biomedical imaging, especially for lesion segmentation (Guerrero et al., 2018; Ghafoorian et al., 2017). For instance, in the MICCAI WMH Segmentation Challenge 2017 (MWSC 2017, https://wmh.isi.uu.nl, Kuijf et al. (2019)), out of 20 methods competing in the initial call, 14 of them (including the top ranking methods) were based on DL (Kuijf et al., 2019). Existing DL methods for WMH segmentation have used convolutional neural network (CNN) models, including various ensemble models (Li et al., 2018; Kuijf et al., 2019), encoder-decoder models (Li et al., 2013; Guerrero et al., 2018; Kuijf et al., 2019; Zhang et al., 2018) - especially U-Nets (proposed by Ronneberger et al. (2015)), 3D multi-dimensional gated recurrent networks (Andermatt et al., 2016) and ResNets (Guerrero et al., 2018). The common choice of inputs used for these networks have been 2D slices (Li et al., 2018) or small 2D/3D patches at multiple scales (Ghafoorian et al., 2017; Andermatt et al., 2016). The choice of the model dimension is affected by various factors such as size and distribution of lesions and amount of data available. Various architectures of 3D CNN models have been successfully used in the segmentation of various types of larger lesions (Kamnitsas et al., 2017; Havaei et al., 2017; Oktay et al., 2018). However, images with WMHs include small lesions in the deep regions, with low contrast and poor context, often with poor resolution along *z*-dimension. This leads to inconsistencies in the detection of WMH boundary voxels using 3D CNNs for lesions of various sizes. From the implementation point of view, 2D models use far fewer parameters when compared to 3D models. However, using 2D models can cause discontinuities in segmentation across the *z*-dimension since individual slices are considered separately. Also for this class of methods, not many are publicly available. In fact, even if most of the pre-trained models from MWSC 2017 challenge are publicly available for testing, there are no independent DL tools available that allow users to train/fine-tune the models on their own data for improving segmentation performance on various datasets from different scanners/centres.

We aimed to develop a DL tool that provides highly accurate segmentation in both periventricular and deep regions, that is publicly available, and that has the flexibility to change training hyperparameters options on various datasets. In this work, we propose TrUE-Net (Triplanar U-Net ensemble network), a DL method for segmentation of WMHs, consisting of an ensemble of U-Nets, each applied to three (axial, sagittal and coronal) planes of structural brain MR images. We aim to improve WMH segmentation irrespective of lesion location by training the TrUE-Net using loss functions that take into account the anatomical location and distribution of WMHs. We evaluate the proposed model on 5 different datasets with different acquisition (scanner and MRI protocol) and lesion characteristics: one from a study on neurodegeneration in prodromal and manifest Alzheimer’s disease, one from a vascular cohort, and three from a publicly available training dataset from MWSC 2017. We compared the performance of TrUE-Net against methods using hand-crafted features and DL methods. Initially, we performed a direct comparison between the results of TrUE-Net and BIANCA, the existing FSL (FMRIB software library) WMH segmentation tool Griffanti et al. (2016), with respect to manual segmentations on the above datasets. Later, we compared the TrUE-Net results with those of the top performing method (Li et al., 2018) from the MWSC 2017 on the publicly available challenge training dataset (Kuijf et al., 2019). Finally, we performed indirect comparisons of TrUE-Net results against various existing methods.

## 2. Materials and methods

### 2.1. Triplanar U-Net Ensemble Network (TrUE-Net)

#### 2.1.1. Preprocessing

We used both T1-weighted and FLAIR images as inputs for the model. We reoriented the images to the standard MNI space, performed skull-stripping FSL BET (Smith, 2002) and bias field correction using FSL FAST (Zhang et al., 2001). We registered the T1-weighted image to the FLAIR using linear rigid-body registration (Jenkinson and Smith, 2001) and cropped the field of vision (FOV) close to the brain and applied Gaussian normalisation to normalise the intensity values. We then extracted 2D slices from the volumes from all three planes. For the axial plane, we cropped the slices to a dimension of 128 × 192 voxels. For sagittal and coronal slices, we cropped and resized the extracted slices to 192 × 120 and 128 × 80 voxels respectively, using bilinear interpolation.

#### 2.1.2. TrUE-Net architecture

The proposed triplanar architecture consists of three 2D networks, each one detecting WMHs from a different plane. The triplanar network reduces discontinuities in WMH segmentation across slices and provides better and comprehensive lesion boundary delineation, using fewer parameters compared to a 3D CNN.

In TrUE-Net, we combined three 2D U-Nets in parallel within an ensemble model. In the ensemble architecture, variation in the individual probability maps (due to noise or spurious structure) is reduced when they are combined in the ensemble network.

Fig. 1 shows the architecture of the proposed TrUE-Net. For each plane, the 2D model takes FLAIR and T1-weighted slices as input channels and provides the probability map in the corresponding plane. In each plane, we trimmed the depth of the classic U-Net (Ronneberger et al., 2015) to obtain a 3-layer deep U-Net model. This reduces the computational load and improves the model sensitivity towards small lesions. Our model mostly uses 3 × 3 convolutional kernels, except for the initial 5 × 5 convolutional kernels in the first layer of the sagittal and coronal U-Nets (figure 1(a)), since larger receptive fields could aid in learning more generic lesion patterns in these planes, thus reducing discontinuities across slices. Each convolution layer is followed by a batch normalisation layer and an activation layer (using *ReLU* - rectified linear unit). We added a 1 × 1 convolutional kernel at the end, before the softmax layer for predicting the probability maps. In the ensemble model, training of U-Nets in the individual planes occurs independently, using the slices extracted from the corresponding planes from the resized training images. During testing, for each network we assembled the slices into a 3D probability map. We then resized each 3D map back to the original dimension and finally averaged the three 3D maps to obtain the final probability map. We then obtained the final probability map by averaging the individual 3D probability maps from the 3 planar orientations, after resizing them back to their original dimensions.

**Figure 1:**
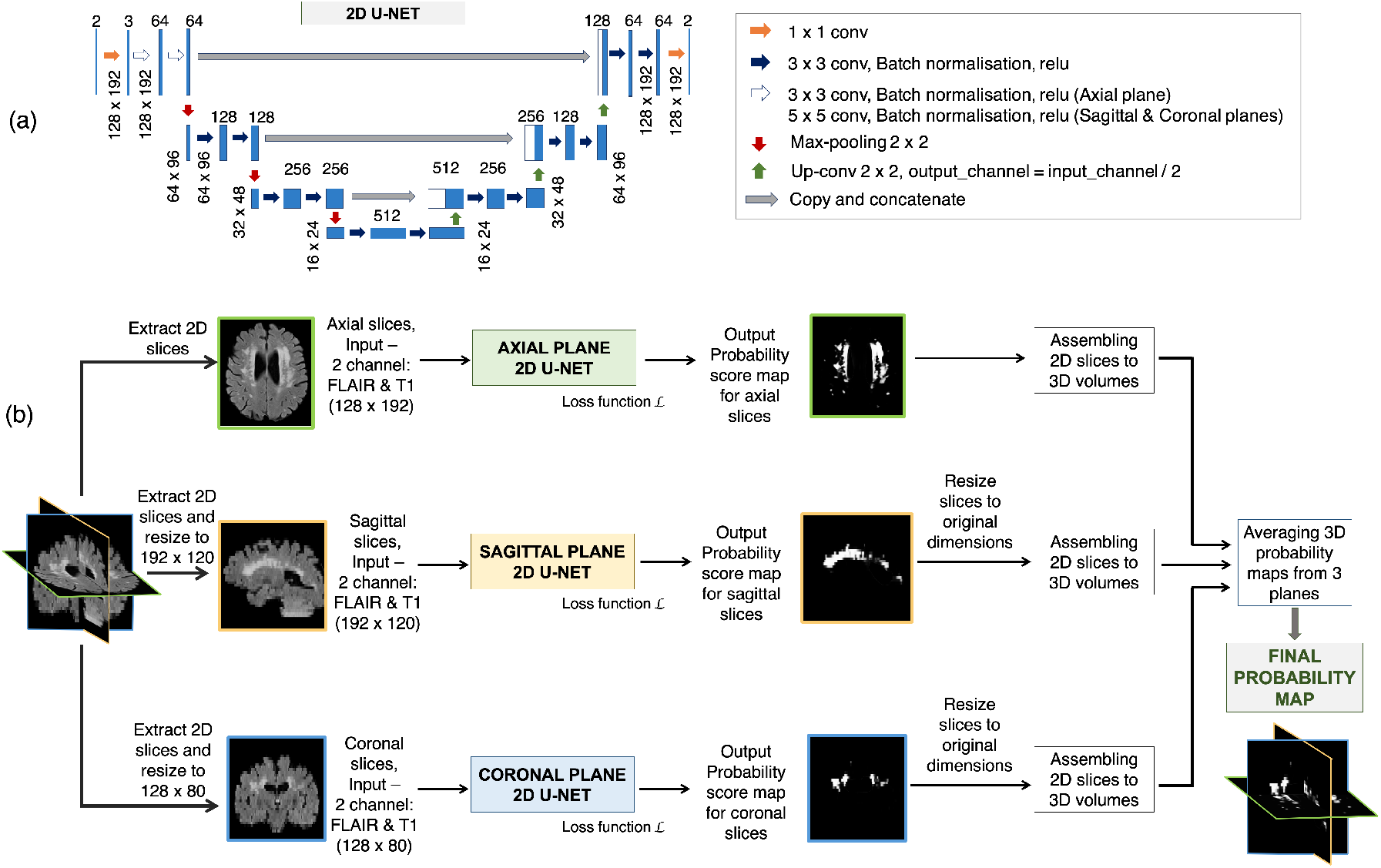
Triplanar U-Net ensemble network (TrUE-Net). (a) U-Net model used in individual planes, (b) Overall TrUE-Net architecture.

#### 2.1.3. Loss function

We used a weighted sum of the voxel-wise cross-entropy (CE) loss function and the Dice loss (DcL) as the total cost function. This is because the CE loss aims to make the segmentation better at the image-level and is biased towards the detection of larger periventricular WMHs, due to the imbalance between WMH and background class samples, especially in deep regions. Hence, we weighted the CE loss function using a spatial weight map (a sample shown in Fig. 2) to up-weight the areas that are more likely to contain the less represented class (i.e. WMH). In addition, we added Dice loss which has stronger sensitivity to missing small WMHs, thereby directly improving the Dice similarity measure of the segmentation. Dice loss has also been shown to work well with unbalanced classes (Milletari et al., 2016; Li et al., 2018).

**Figure 2:**
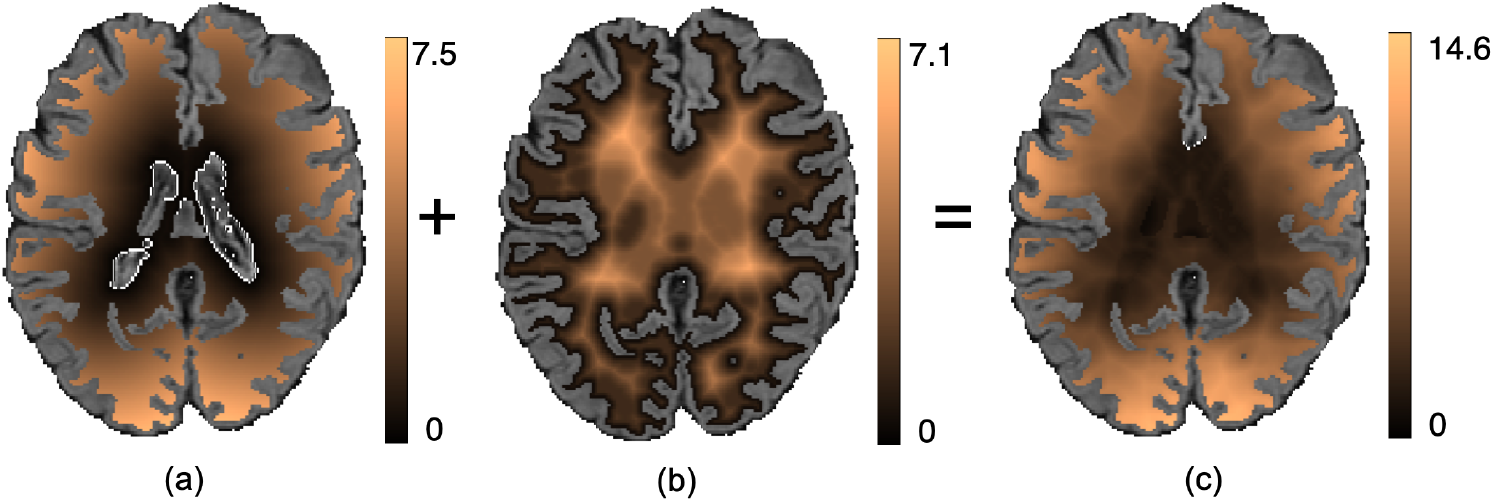
Example weight maps for weighting the voxel-wise cross-entropy loss function. Maps showing the sum of (a) Distance from ventricles *D*_*vent*_ and (b) Distance from gray matter (GM) *D*_*GM*_ to get (b) Final weight map *D*_*wei*_.

We determined the distance from ventricles, 0 ≤ *D*_*vent*_ ≤ *N*_*D*_, *N*_*D*_ ∈ *R* (Fig. 2a), and the distance from the brain gray matter (GM), 0 ≤ *D*_*GM*_ ≤ *M*_*D*_, *M*_*D*_ ∈ *R* (Fig. 2b) and obtained the final weight map as the sum of both, *D*_*wei*_ = *D*_*vent*_ + *D*_*GM*_, 0 ≤ *D*_*wei*_ ≤ (*N*_*D*_ + *M*_*D*_) (Fig. 2c), where *N*_*D*_ and *M*_*D*_ are maximum distances from ventricles and GM respectively (all distances in mm). Hence the total weighted CE loss for *N* voxels is given by,

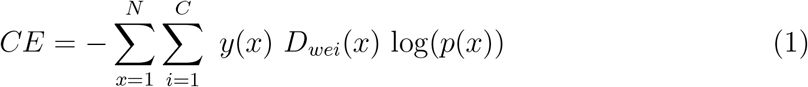

where *p*(*x*) denotes the output of the soft-max layer, *C* is the number of classes and *y*(*x*) = 0 or 1, the value at each voxel *x* on the manual segmentation. Given the manual segmentation *M* and binary map *P*_*th*_ obtained by thresholding predicted probability maps, the Dice loss for *N* voxels is given by,

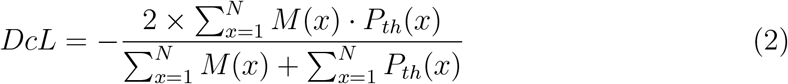

Hence the total loss function *L* is given by,

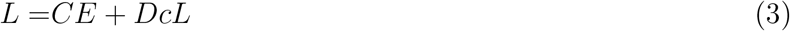

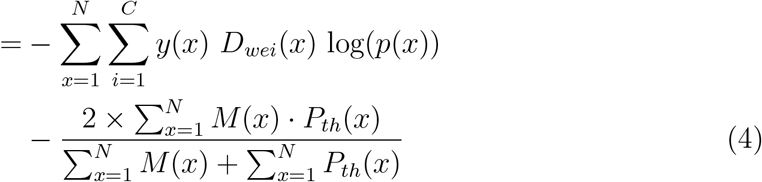

We chose equal weights on the basis of initial empirical experiments (not shown) for the Dice loss and the weighted CE loss while adding them to obtain the total loss.

#### 2.1.4. Post-processing

We padded the output probability maps with zeros to bring them back to their original dimensions. We later masked the probability map with the white matter mask obtained from a dilated and inverted cortical CSF tissue segmentation combined with other deep grey exclusion masks (using FSL FAST (Zhang et al., 2001) and *make bianca mask* command in FSL BIANCA (Griffanti et al., 2016)). Finally, we thresholded the masked probability maps at 0.5.

#### 2.1.5. Implementation details

We implemented the network in Python 3.6 using Torch 1.2.0. The network was trained on an NVIDIA Tesla V100, taking 80 epochs at 45 seconds (for 3 planes) per epoch for 15,000 samples with the training/validation split of 90%/10%. We used the Adam Optimiser with *ϵ*=10^−4^. We used a batch size of 8, with an initial learning rate of 1×10^−3^ and reducing it by a factor 1×10^−1^ every 2 epochs, until it reaches 1×10^−5^, after which we maintain the fixed learning rate value. We chose the above parameters empirically based on the model convergence (refer to section 2.3.2 for more details). Data augmentation was applied using translation (x/y-offset ∈ [-10, 10]), rotation (*θ* ∈ [-10, 10]), random noise injection (Gaussian, *μ* = 0, *σ*^2^ ∈ [0.01, 0.09]) and Gaussian filtering (*σ* ∈ [0.1, 0.3]), increasing the dataset by a factor of 10 and 6 for axial and sagittal/coronal planes respectively. The hyperparameter values for the data augmentation transformations were randomly sampled from the closed intervals specified above using a uniform distribution.

### 2.2. Datasets

#### 2.2.1. Neurodegenerative cohort (NDGEN)

The dataset, used in Zamboni et al. (2013), includes MRI data from 9 subjects with probable Alzheimer’s Disease, 5 with amnestic mild cognitive impairment and 7 cognitively healthy control subjects (age range 63 - 86 years; mean age 77.1 ± 5.8 years; median age 77 years; F:M = 10:11). The total brain volume range: 1189282 - 1614799 mm^3^, median: 1424669 mm^3^. Manual segmentation was available for all datasets (WMH load range: 1878 - 89259 mm^3^, median: 20772 mm^3^). The images were acquired using a 3T Siemens Trio Scanner, with FLAIR (TR/TE = 9000/89 ms, flip angle 150°, FOV 220 mm, voxel size 1.1 × 0.9 × 3 mm, matrix size 256 × 256 × 35 voxels) and T1-weighted sequence (3D MP-RAGE sequence, TR/TE = 2040/4.7 ms, flip angle 8°, FOV 192 mm, voxel size 1 mm isotropic, matrix size 174 × 192 × 192 voxels).

#### 2.2.2. Vascular cohort - Oxford Vascular Study (OXVASC)

The dataset consists of 18 participants in the OXVASC study (Rothwell et al., 2004), who had recently experienced a minor non-disabling stroke or transient ischemic attack (age range 50 - 91 years; mean age 73.27 ± 12.32 years; median age 75.5 years; F:M = 7:11). The total brain volume: range: 1290926 - 1918604 mm^3^, median: 1568233 mm^3^. Manual segmentation was available for all datasets (WMH load range: 3530 - 83391 mm^3^, median: 16906 mm^3^). The images were acquired using a 3T Siemens Trio Scanner, with FLAIR (TR/TE = 9000/88 ms, flip angle 150°, voxel size 1 × 3 × 1 mm, matrix size 174 × 52 × 192 voxels) and T1-weighted sequence (3D MP-RAGE sequence, TR/TE = 2000/1.94 ms, flip angle 8°, voxel size 1 mm isotropic, matrix size 208 × 256 × 256 voxels).

#### 2.2.3. MICCAI WMH Segmentation Challenge Dataset (MWSC)

The dataset consists of 60 subjects from three different sources (20 subjects each) provided as training sets for the challenge (http://wmh.isi.uu.nl/): UMC Utrecht, NUHS Singapore and VU Amsterdam. The brain volume ranges are 1257820 - 1844920 mm^3^ (median – 1473389 mm^3^) for UMC Utrecht, 1147248 - 1532268 mm^3^ (median: 1351325 mm^3^) for NUHS Singapore and 1219614 - 1787321 mm^3^ (median: 1441201 mm^3^) for VU Amsterdam. Manual segmentations are available for all three datasets, with an additional exclusion label provided for other pathologies. In the challenge, the masks with exclusion labels were ignored during performance evaluation. However, we included these masks as parts of non-lesion tissue, during both training and testing, for the calculation of the performance metrics in order to get more stringent evaluation in the presence of other pathologies. The WMH volume ranges (excluding other pathologies) are 845 - 74991 mm^3^ (median: 26240 mm^3^) for UMC Utrecht, 786 - 61332 mm^3^ (median: 17795 mm^3^) for NUHS Singapore and 1522 - 43528 mm^3^ (median: 6015 mm^3^) for VU Amsterdam. For more details regarding MRI acquisition parameters, refer to http://wmh.isi.uu.nl/.

### 2.3. Experiments

#### 2.3.1. Performance evaluation metrics

We used the following performance metrics in our evaluation:

- **Dice Similarity Index (SI)**, calculated as 2 × (true positive WMH voxels) / (true WMH voxels + positive WMH voxels).
- **Voxel-wise true positive rate (TPR)** is the ratio of the number of true positive WMH voxels to the number of true WMH voxels.
- **Voxel-wise false positive rate (FPR)** is the number of false positive (FP) WMH voxels divided by the number of non-WMH voxels.
- **Cluster-wise TPR** is the number of true positive WMH clusters divided by the total number of true WMH clusters.
- **Absolute volume difference (AVD) (%)** is the absolute difference between the volume of manually segmented WMHs voxels and the volume of detected WMHs voxels, as a percentage of the manually segmented lesion voxels.
- 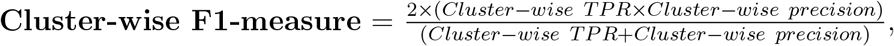, where Cluster-wise precision is the number of true positive WMH clusters divided by the total number of detected WMH clusters.
- **95^th^ percentile of Hausdorff distance measure (H95)**: We determined the set of the closest distances between the points on the detected WMH boundary and the manually segmented WMH boundary. We calculated the 95^th^ percentile of this set of closest distance values, instead of the maximum value used for the standard Hausdorff value.

#### 2.3.2. Effect of training hyperparameters on model performance

We used the NDGEN dataset for the initial optimisation of network parameters, and for determining the number of epochs. We explored the effect of batch-size, learning rate and epsilon (*ϵ*) value of the Adam optimiser on model convergence. Batch size is the number of training data used by the optimiser at a given instance for updating model parameters. The parameter *ϵ* is used in the denominator of the weight update function in the optimiser to avoid the divide-by-zero error. Learning rate determines the extent of change/adjustment of weights with respect to the gradient. We experimented with three batch sizes: 8, 16 and 32, and three *ϵ* values: 1 × 10^−2^, 1 × 10^−4^ and 1 × 10^−6^. For learning rate, we tested the effect of following 3 settings: (i) **Higher:** Initially 1 × 10^−2^, reducing by a factor 1 × 10^−1^ every 2 epochs until it reaches 1 × 10^−4^ and keeping it constant afterwards, (ii) **Medium:** Initially 1 × 10^−3^, reducing by a factor 1 × 10^−1^ every 2 epochs until it reaches 1 × 10^−5^ and (iii) **Lower:** Initially 1 × 10^−4^, reducing by a factor 1 × 10^−1^ every 2 epochs until it reaches 1 × 10^−6^.

#### 2.3.3. Ablation study: effect of loss function components on the segmentation performance

We performed an ablation study to determine the effect of the components of the loss function on the segmentation results using the NDGEN dataset. For this experiment, we removed one component of the loss function at a time and compared the performance of TrUE-Net with three cases of loss functions: cross-entropy loss, weighted cross-entropy loss and weighted cross-entropy loss + Dice loss. We maintained the same training and model parameters for all three options. Since we hypothesized that the weighting of the CE loss would specifically improve the performance in deep regions, we determined SI values in periventricular and deep regions separately, and performed paired t-tests to compare the results between regions. We adopted the 10 mm distance rule (DeCarli et al., 2005; Griffanti et al., 2018) for classification of PWMH and DWMH.

#### 2.3.4. Leave-one-out evaluation of WMH segmentation

We performed leave-one-out (LOO) evaluation of our proposed model on all the subjects from the MWSC dataset (combined from Utrecht, Singapore and Amsterdam cohorts), NDGEN and OXVASC datasets, based on the performance metrics specified in section 2.3.1. We also determined the performance of the model in the deep and the periventricular regions (using 10 mm distance rule as above) separately, and performed paired t-tests to compare the results between regions.

#### 2.3.5. Comparison with BIANCA

We performed a direct comparison of our TrUE-Net with BIANCA using LOO evaluation on the MWSC, NDGEN and OXVASC datasets, based on the performance metrics specified in section 2.3.1 and performed paired t-tests between TrUE-Net and BIANCA results.

##### BIANCA features and training options

For NDGEN, we used FLAIR and T1 as features and the options that provided the best results in the initial validation of BIANCA Griffanti et al. (2016). Other than the default options, we used the following non-default options: location of training points = no border, number of training points = Fixed + unbalanced with 2000 lesion points and 10,000 non-lesion points. The ‘no border’ option and fixed + unbalanced lesion and non-lesion points have been shown to provide the best results during initial validation of BIANCA (Griffanti et al., 2016) and also in our initial tests on the same data. For OXVASC we used FLAIR + T1 + MD as features. Specific options used were: sw = 2, 3D patch with patch size of 3. Due to the anisotropic nature of the voxels, additional intensity features obtained by averaging over a smaller 3D patch provided better results during the initial tests. For the MWSC dataset, we trained BIANCA using the same features and BIANCA options used for NDGEN. For all the datasets, we applied a global threshold value of 0.9 on the BIANCA lesion probability maps and masked with the white matter mask (obtained in section 2.1.4) to obtain the final binary lesion maps.

#### 2.3.6. Comparison with the top ranking method of MWSC 2017

We compared the LOO results of TrUE-Net with those reported in the supplementary table S1 in Li et al. (2018), which is the top ranking method in MWSC 2017 (Kuijf et al., 2019) on the 60 subjects from the MWSC cohorts. We also performed paired t-tests between the performance measure of TrUE-Net and Li et al. (2018).

#### 2.3.7. Comparison with other existing methods

Finally, we performed an indirect comparison of our results with those obtained by other WMH segmentation methods in the literature. We included in our comparisons only methods that achieved minimum Dice overlap metric or voxel-wise TPR of 0.70.

## 3. Results

### 3.1. Effect of training parameters on model performance

Figure 3 shows the loss function decay with different batch sizes, learning rates and epsilon (*ϵ*) values. In all the cases, the model started to converge at approximately 80 epochs.

**Figure 3:**
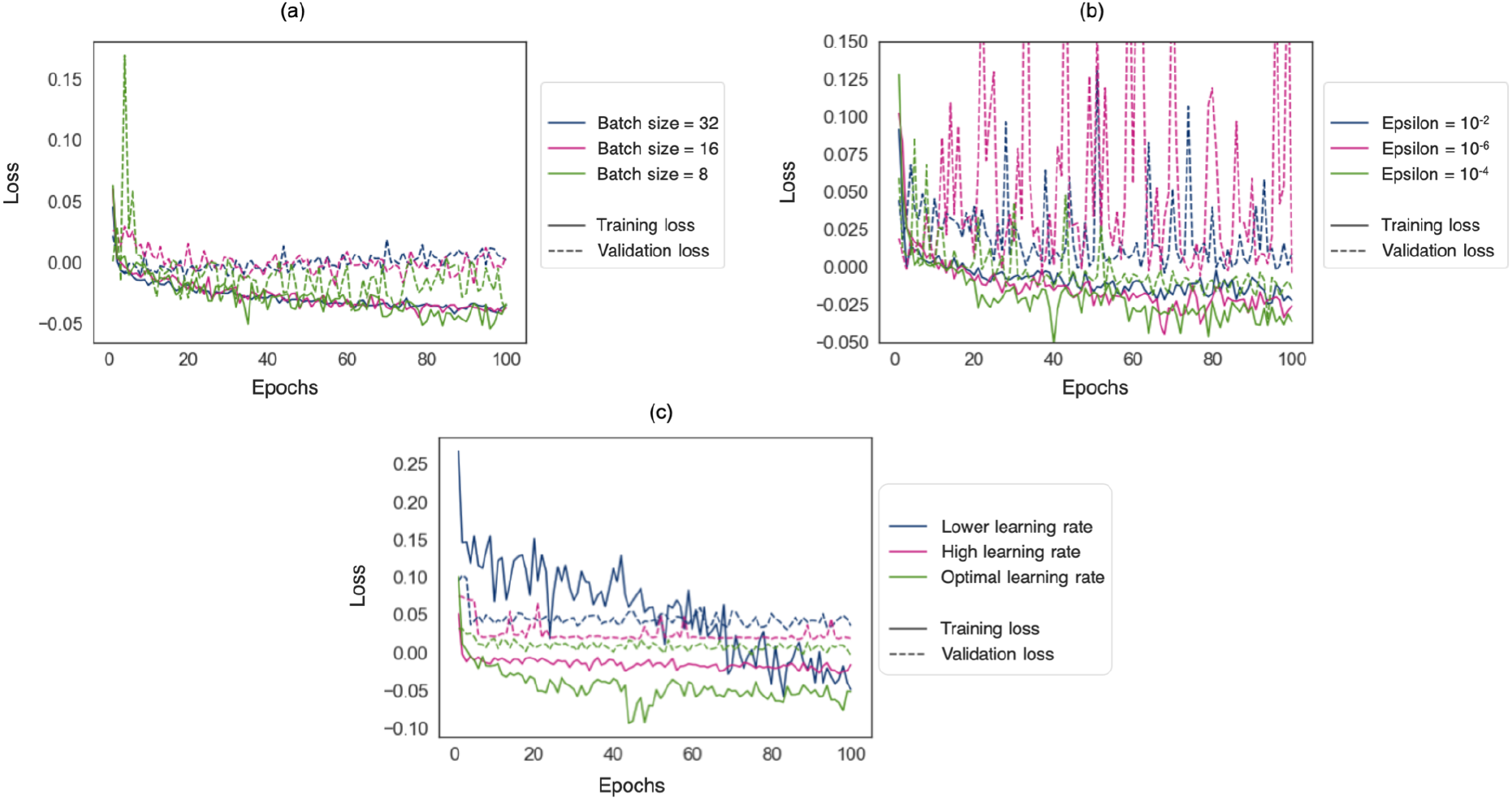
Effect of training parameters on model convergence on the NDGEN dataset. Training and validation loss decays have been shown for different (a) batch sizes, (b) epsilon (*ϵ*) values and (c) learning rates. The plots shown in green correspond to the optimal values chosen for each parameter.

At a batch size of 8, the loss function became noisier, but had lower values. Larger batch sizes resulted in over-fitting, as evident from higher validation loss values for batch sizes of 16 and 32 (figure 3a). Therefore, we chose a batch size of 8 for our experiments henceforth. As the *ϵ* value gets smaller in the denominator, the optimiser makes larger weight updates leading to unstable optimisation, as shown in figure 3b for an *ϵ* value of 1 × 10^−6^. Since an *ϵ* value of 1 × 10^−2^ provided higher loss values (with slightly unstable validation loss), we set the *ϵ* value to the optimal value of 1 × 10^−4^ for all the subsequent experiments. From figure 3c, for a lower learning rate, the loss decay was slower and hence required more epochs for convergence. On the other hand, for a higher learning rate the loss values values converged before 20 epochs, although with higher loss values. At the optimal learning, the loss decay was slow and converged to a much lower loss value for both training and validation datasets. Hence, we chose the optimal learning rate schedule to be from 1 × 10^−3^ to 1 × 10^−5^.

### 3.2. Ablation study: effect of loss function components on segmentation performance

The boxplots of the validation SI values for WMH segmentation in deep and periventricular areas are shown in Figure 4. On performing a one-way ANOVA across three components of the loss function (CE loss, weighted CE loss and weighted CE + Dice loss), we found that the SI values significantly increase (F = 25.39, p < 0.0001) for DWMHs with the addition of each component of the loss function. Both weighted CE loss and weighted CE + Dice loss gives significantly higher SI values than the CE loss component alone in the deep regions, whereas in the periventricular regions the SI values were significantly higher after adding the Dice loss component.

**Figure 4:**
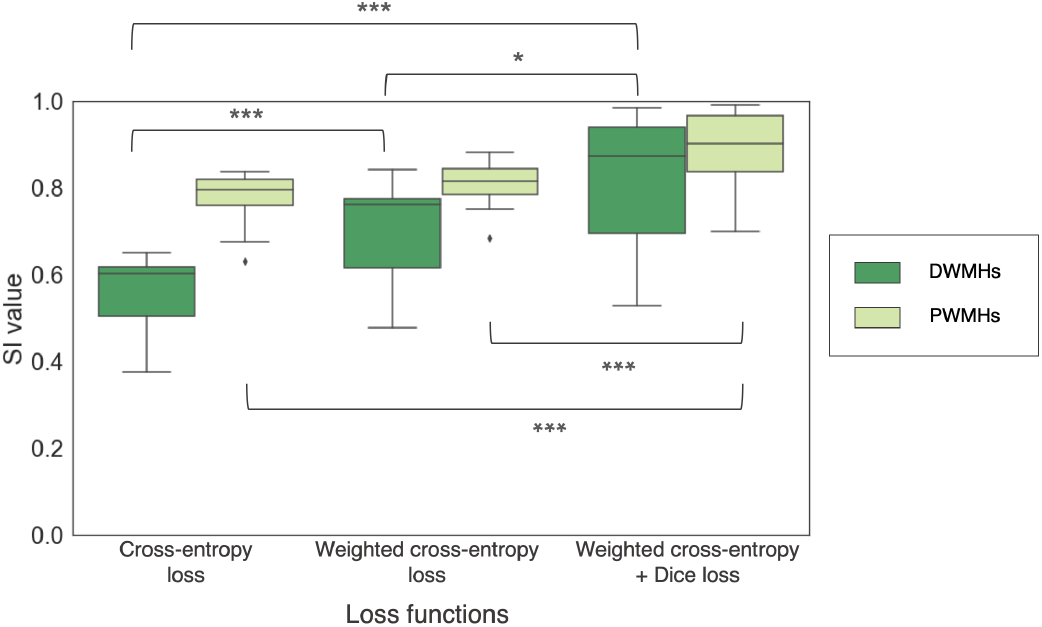
Boxplots of SI values obtained for deep and periventricular regions on the NDGEN dataset for CE, weighted CE and weighted CE + Dice loss functions. *** - p < 0.0001, ** - p < 0.001, * - p < 0.01

Figure 5 shows the effect of CE and Dice loss components on TrUE-Net segmentation for a subject with medium lesion load. From the boxplots and the visual results, the effect of the composition of the loss function was observable mainly along the edges of PWMHs and DWMHs. In general, we have a large imbalance between WMH and normal WM classes and aim to focus on detecting more true WMHs. Since the CE loss function aggregates the loss values at individual voxels into an image-level value, it is generally biased towards WMHs in periventricular regions (figure 5c) where the imbalance between WMH and non-WMH voxels is less. Therefore, in some cases, the segmentation results extended beyond the ventricle lining. Using weighted CE loss controls this behaviour, since it relies on the weight maps (described in section 2.1.3) prepared with prior anatomical information. Overcoming this class imbalance, DWMHs are given weights that are more similar to those given to PWMHs. As a result, more DWMHs are detected and the over-segmentation in periventricular regions is avoided (figure 5d). Since missing small WMHs would make more difference to the Dice component than to the CE loss component, our expectation would be that training the network with the inclusion of a Dice component in the loss function would favour finding small lesions and reducing false negatives (especially in the areas of high class imbalance). Therefore, combining the advantages of both, the Dice loss and the weighted CE loss components, provided better detection of DWMHs, along with accurate segmentation of PWMH borders (figure 5). The effect was particularly observable in low lesion load subjects, where the addition of the Dice loss component provided more precise segmentation, with fewer false positives. Figure 6 shows the effect of lesion load on SI values of TrUE-Net segmentation for the three loss function compositions. In all three cases there was a significant correlation between SI and lesion load (CE loss: *ρ* = 0.48, p=0.03; weighted CE loss: *ρ* = 0.62, p=0.003; weighted CE + Dice loss: *ρ* = 0.40, p=0.08). The combination of the weighted CE and Dice loss components achieved higher SI values and this case was less affected by lesion load, as indicated by the slightly lower correlation value. This correlation between SI values and lesion load reflects a variation in performance with the amount of WMHs that are present and is undesirable for robust performance. However, the correlation coefficients of the cases with the CE loss component and weighted CE loss component were not significantly higher than the combination case (p = 0.41 and p = 0.78 respectively).

**Figure 5:**
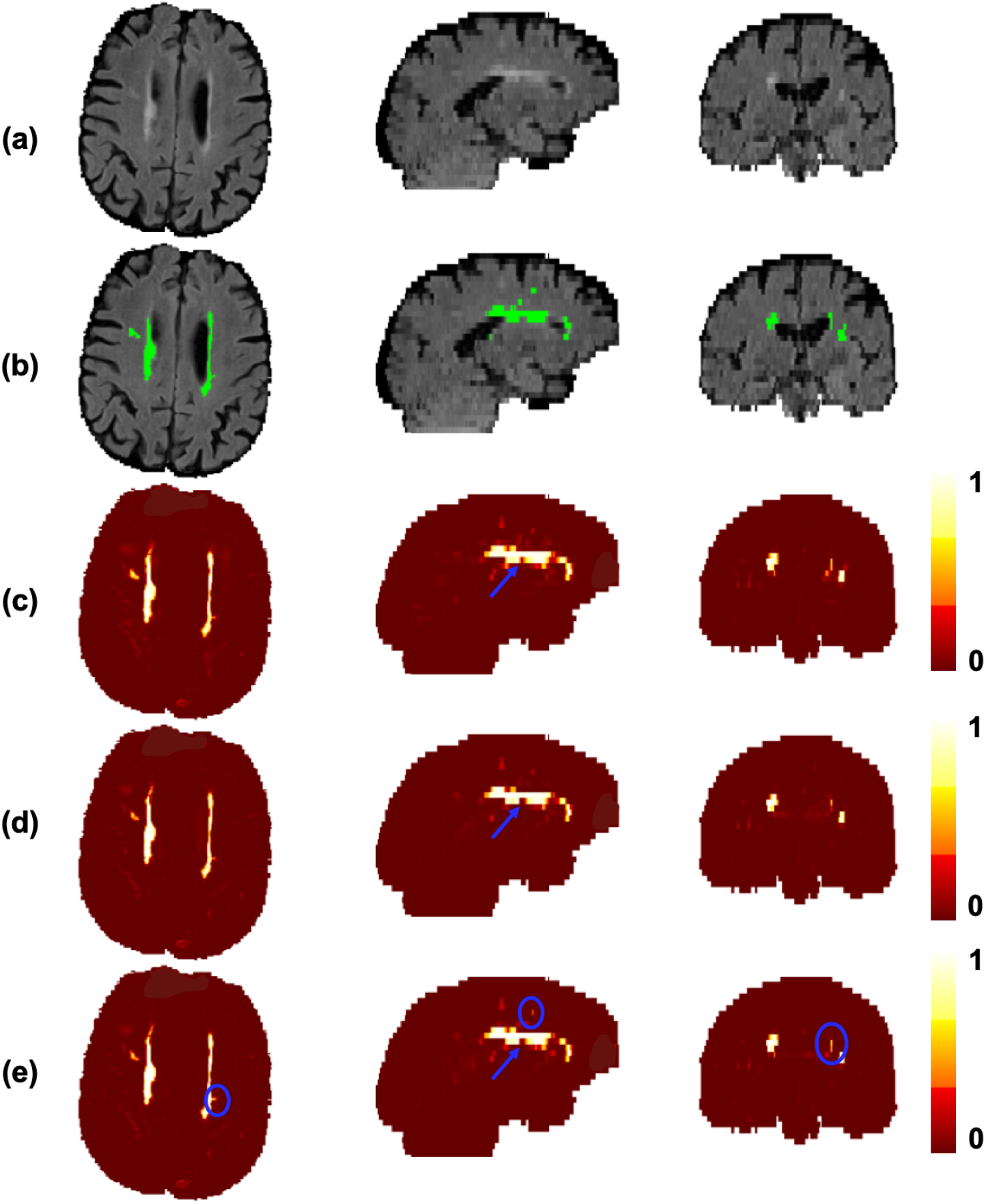
Effect of loss functions on the segmentation performance in the NDGEN dataset. An example showing (a) FLAIR images, (b) manual segmentation against the results of (c) cross-entropy (CE) loss, (d) weighted cross-entropy loss and (e) weighted cross-entropy loss + Dice loss. Differences in lesion segmentations indicated by blue circles and arrows.

**Figure 6:**
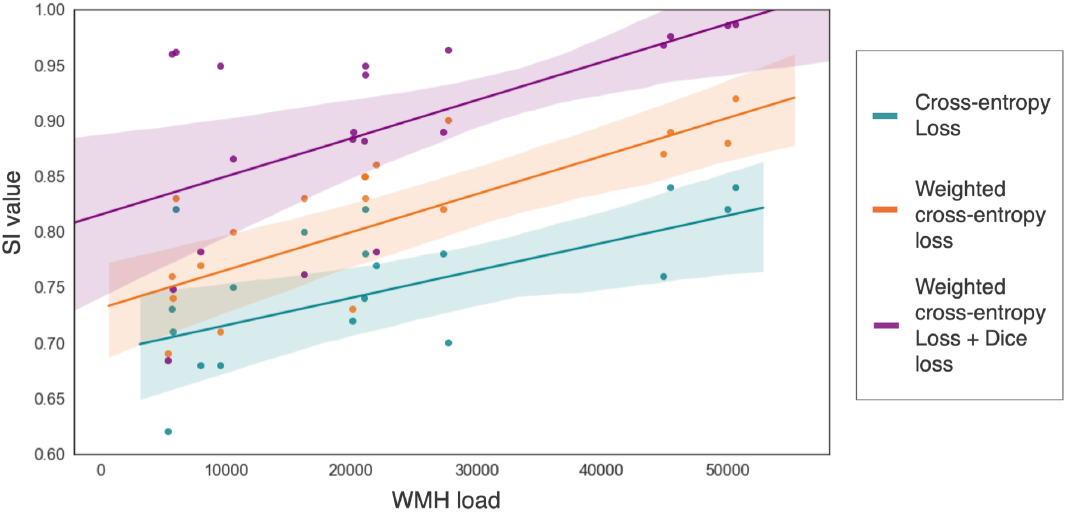
Regression plot showing the impact of lesion load on SI values for weighted cross-entropy loss (magenta) and weighted cross-entropy loss + Dice loss (blue), on the NDGEN dataset. The shaded region represents the 95% confidence interval of the regression estimates.

### 3.3. Evaluation of WMH segmentation performance

Figure 7 shows the boxplots of the performance metrics for the leave-one-out (LOO) validation on the MWSC cohorts, NDGEN and OXVASC datasets. Table 1 shows the corresponding values. TrUE-Net achieved its best performance on the MWSC and OXVASC datasets. Within the MWSC cohorts, TrUE-Net achieved the best performance for the Utrecht cohort (SI: 0.92 ± 0.04, voxel-wise TPR: 0.90 ± 0.07, voxel-wise FPR: 0.7 × 10^−4^, cluster-wise TPR: 0.88 ± 0.08, cluster-wise F1-measure: 0.92 ± 0.07, AVD: 10.95 ± 6.6%, H95: 1.25 ± 0.72 mm). The model achieved the lowest performance on the NDGEN dataset, however with the highest cluster-wise TPR value of 0.87 ± 0.12. This indicates that it detects more true positive lesions when compared to other datasets, while the lesion boundaries were better delineated in other datasets.

**Figure 7:**
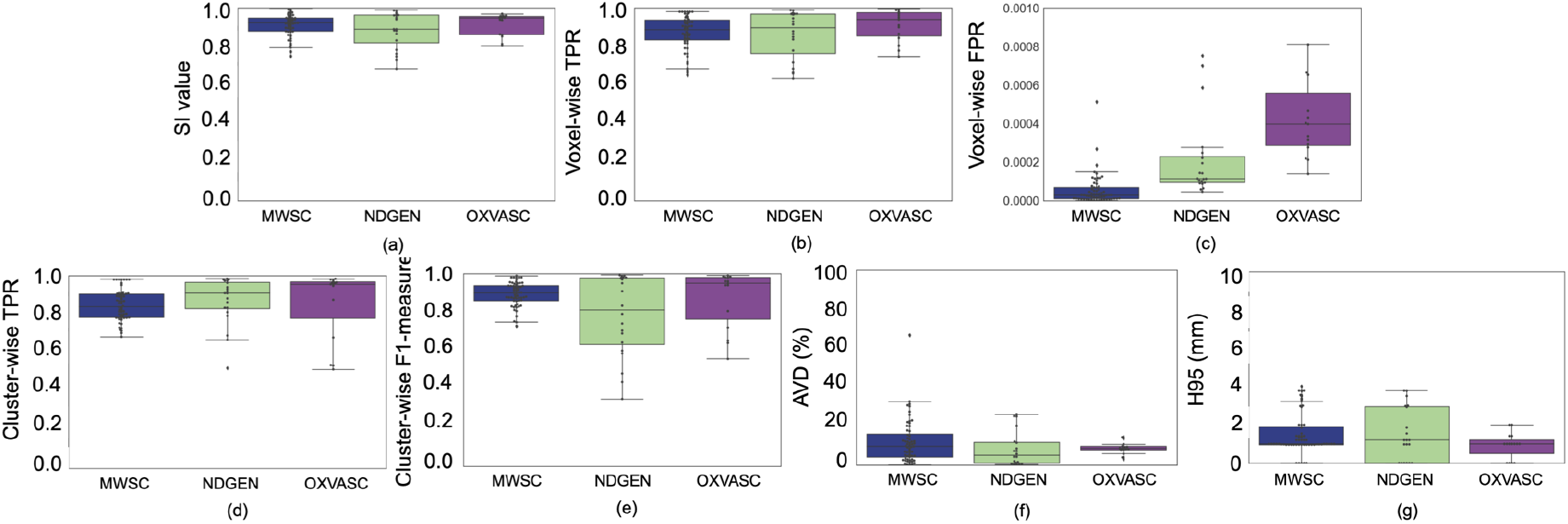
Boxplots of performance metrics obtained from LOO evaluation on the MWSC, NDGEN and OXVASC datasets - (a) SI value, (b) voxel-wise TPR, (c) voxel-wise FPR, (d) cluster-wise TPR, (e) cluster-wise F1-measure, (f) Absolute volume difference (AVD), and (g) 95^th^ percentile of Hausdorff distance.

**Table 1:** LOO evaluation of TrUE-Net on the MWSC, NDGEN and OXVASC datasets (the best value for each measure across the datasets is highlighted in bold).

| Perf. metrics | MWSC | NDGEN | OXVASC |
| --- | --- | --- | --- |
| SI | **0.91 ± 0.06** | 0.88 ± 0.1 | **0.91 ± 0.06** |
| Voxel-wise TPR | 0.87 ± 0.08 | 0.86 ± 0.12 | **0.91 ± 0.08** |
| Voxel-wise FPR | **0.5 × 10<sup>-4</sup>** | 2.1 × 10<sup>-4</sup> | 4.4 × 10<sup>-4</sup> |
| Cluster-wise TPR | 0.84 ± 0.08 | **0.87 ± 0.12** | 0.85 ± 0.18 |
| Cluster-wise F1 measure | **0.89 ± 0.06** | 0.78 ± 0.21 | 0.87 ± 0.15 |
| AVD (%) | 11.6 ± 11.4 | **8.1 ± 9.0** | 8.5 ± 3.0 |
| H95 (mm) | 1.5 ± 1.1 | 1.5 ± 1.4 | **0.9 ± 0.7** |

The visual results of LOO evaluations on the MWSC cohorts, NDGEN and OXVASC datasets are illustrated by a few examples shown in figure 8. In both high and low lesion load cases, TrUE-Net provided an accurate segmentation with respect to manual segmentation without detecting many false positives. In particular, TrUE-Net detected the deep lesions in the low lesion load cases, including subtle ones (figure 8f and j). It is also worth noting that, although manual segmentation is used as gold standard for evaluation, there might be inconsistencies and errors that would affect the performance evaluation. For instance, in a high lesion load example from the Utrecht cohort in figure 8a, some CSF voxels were included were included in the manual segmentation (indicated by a circle), while TrUE-Net successfully excluded them by providing lower probability values in those regions.

**Figure 8:**
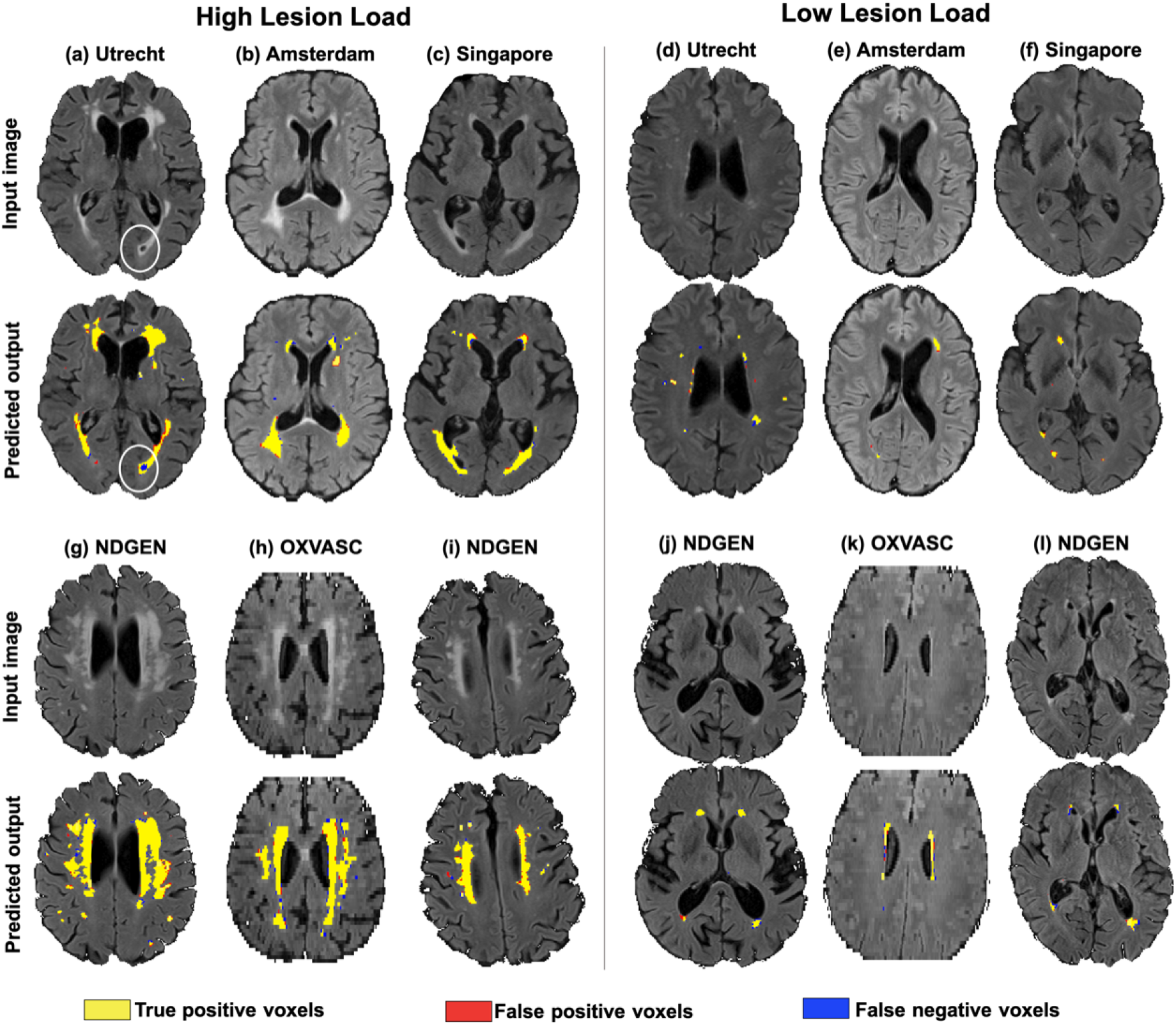
Sample results of TrUE-Net segmentation on the high and low lesion cases - (a, d) Utrecht, (b, e) Amsterdam and (c, f) Singapore cohorts of MWSC dataset, (g, i, j, l) NDGEN and (h, k) OXVASC datasets. True positive, false positive and false negative voxels are indicated in yellow, red and blue respectively.

### 3.3.1. Performance in deep and periventricular lesions

Figure 9 shows boxplots of the performance metrics for DWMHs and PWMHs. Table 2 reports the corresponding descriptive statistics, along with the p-values of the paired t-tests performed between DWMHs and PWMHs. Most of the performance metrics are not significantly different between PWMHs and DWMHs. Particularly, none of the differences in cluster-wise and voxel-wise TPRs are significant, indicating that TrUE-Net not only successfully detects true lesions in both the periventricular and deep regions, but also segments the lesion boundaries accurately in both regions. The only significant differences can be found in the cluster-wise F1-measure in the MWSC dataset, as well as AVD and voxel-wise FPR values in the OXVASC dataset. In the case of the MWSC dataset, the cluster-wise F1-measure in the deep regions were significantly higher than in the periventricular regions. This means that more true DWMHs were detected, with higher cluster-wise precision, compared to PWMHs. In the OXVASC dataset, while DWMHs showed significantly higher AVD percentage values compared to PWMHs, they still correspond to much lower lesion volumes when compared to PWMHs. For instance, AVD value of 25% in the deep region corresponds to a lesion volume of around 600 mm^3^ while the same value corresponds to a much higher value, of around 3800 mm^3^, in the periventricular region.

**Figure 9:**
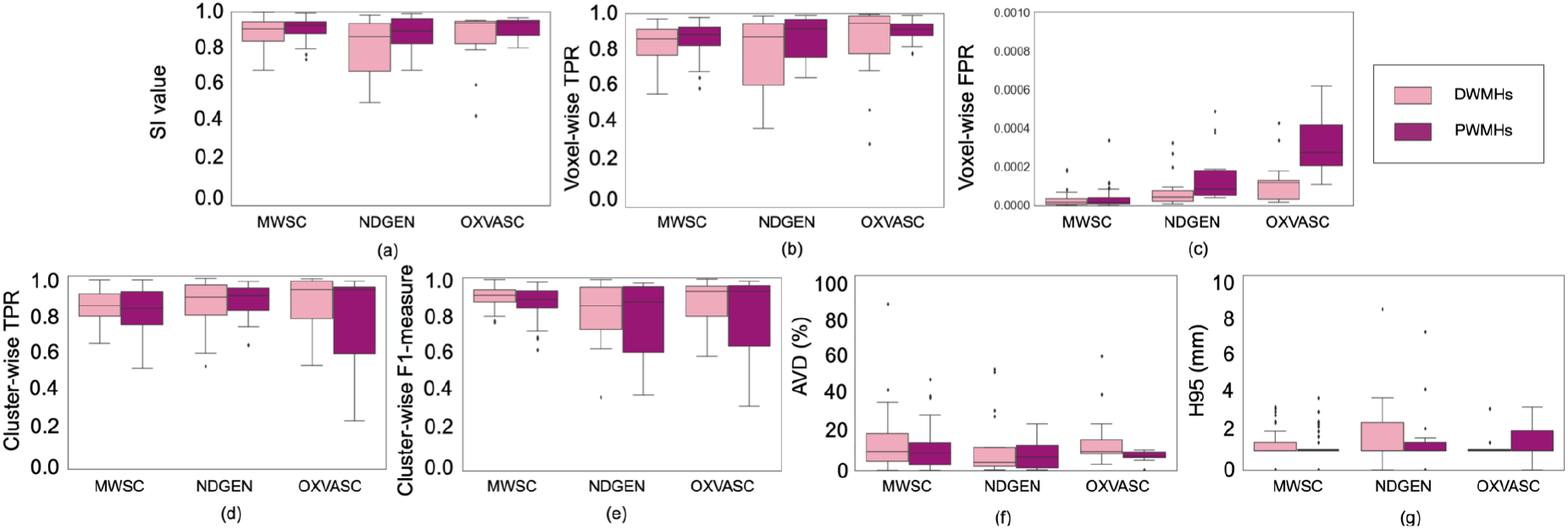
Boxplots of performance metrics obtained in deep and periventricular regions for LOO evaluation on the MWSC, NDGEN and OXVASC datasets - (a) SI value, (b) voxel-wise TPR, (c) voxel-wise FPR, (d) cluster-wise TPR, (e) cluster-wise F1-measure, (f) Absolute volume difference (AVD), and (g) 95^th^ percentile of Hausdorff distance.

**Table 2:** Comparison of performance metrics between PWMHs and DWMHs, along with p-values of two-tailed paired t-test results on the MWSC, NDGEN and OXVASC datasets (the best performance values and significant p-values highlighted in bold).

|  |  | MWSC | NDGEN | OXVASC |
| --- | --- | --- | --- | --- |
| SI | DWMHs | $0.90 \pm 0.07$ | $0.82 \pm 0.15$ | $0.87 \pm 0.15$ |
| | PWMHs | $0.91 \pm 0.06$ | $0.89 \pm 0.09$ | $0.92 \pm 0.05$ |
|  | p-value | 0.07 | 0.07 | 0.21 |
| Voxel-wise<br>TPR | DWMHs | $0.85 \pm 0.10$ | $0.79 \pm 0.12$ | $0.85 \pm 0.20$ |
| | PWMHs | $0.88 \pm 0.08$ | $0.87 \pm 0.12$ | $0.91 \pm 0.06$ |
|  | p-value | 0.07 | 0.13 | 0.32 |
| Voxel-wise<br>FPR | DWMHs | $0.7 \times 10^{-5}$ | $4.1 \times 10^{-5}$ | $4.2 \times 10^{-5}$ |
| | PWMHs | $0.7 \times 10^{-4}$ | $1.2 \times 10^{-4}$ | $3.3 \times 10^{-4}$ |
|  | p-value | 0.45 | 0.07 | **<0.001** |
| Cluster-wise<br>TPR | DWMHs | $0.84 \pm 0.08$ | $0.82 \pm 0.09$ | $0.87 \pm 0.15$ |
| | PWMHs | $0.82 \pm 0.11$ | $0.87 \pm 0.08$ | $0.79 \pm 0.24$ |
|  | p-value | 0.21 | 0.74 | 0.34 |
| Cluster-wise<br>F1-measure | DWMHs | $0.90 \pm 0.05$ | $0.83 \pm 0.20$ | $0.87 \pm 0.12$ |
| | PWMHs | $0.87 \pm 0.08$ | $0.78 \pm 0.20$ | $0.82 \pm 0.21$ |
|  | p-value | **0.03** | 0.39 | 0.40 |
| AVD (%) | DWMHs | $14.0 \pm 14.1$ | $11.3 \pm 8.1$ | $15.9 \pm 14.7$ |
| | PWMHs | $10.9 \pm 10.3$ | $8.17 \pm 8.12$ | $7.5 \pm 2.7$ |
|  | p-value | 0.16 | 0.44 | **0.04** |
| H95 (mm) | DWMHs | $1.2 \pm 0.7$ | $2.4 \pm 1.6$ | $1.2 \pm 0.6$ |
| | PWMHs | $1.5 \pm 1.9$ | $1.4 \pm 1.6$ | $1.4 \pm 1.0$ |
|  | p-value | 0.29 | 0.23 | 0.53 |

Overall, TrUE-Net provided performance metrics for DWMHs on par with PWMHs. This shows that the model provides good delineations of WMHs, with good sensitivity and specificity in both regions.

### 3.4. Comparison with BIANCA

Figure 10 illustrates a few example segmentations obtained with TrUE-Net and BIANCA, with respect to manual segmentation, in the order of decreasing lesion load. TrUE-Net provided more accurate segmentations than BIANCA, especially in the low lesion load subjects. From the figure it can be observed that, as the lesion load decreases, BIANCA detected more false positives, particularly around the ventricles (shown in figure 10a, d and e) and oversegmented PWMHs (figure 10b and c).

**Figure 10:**
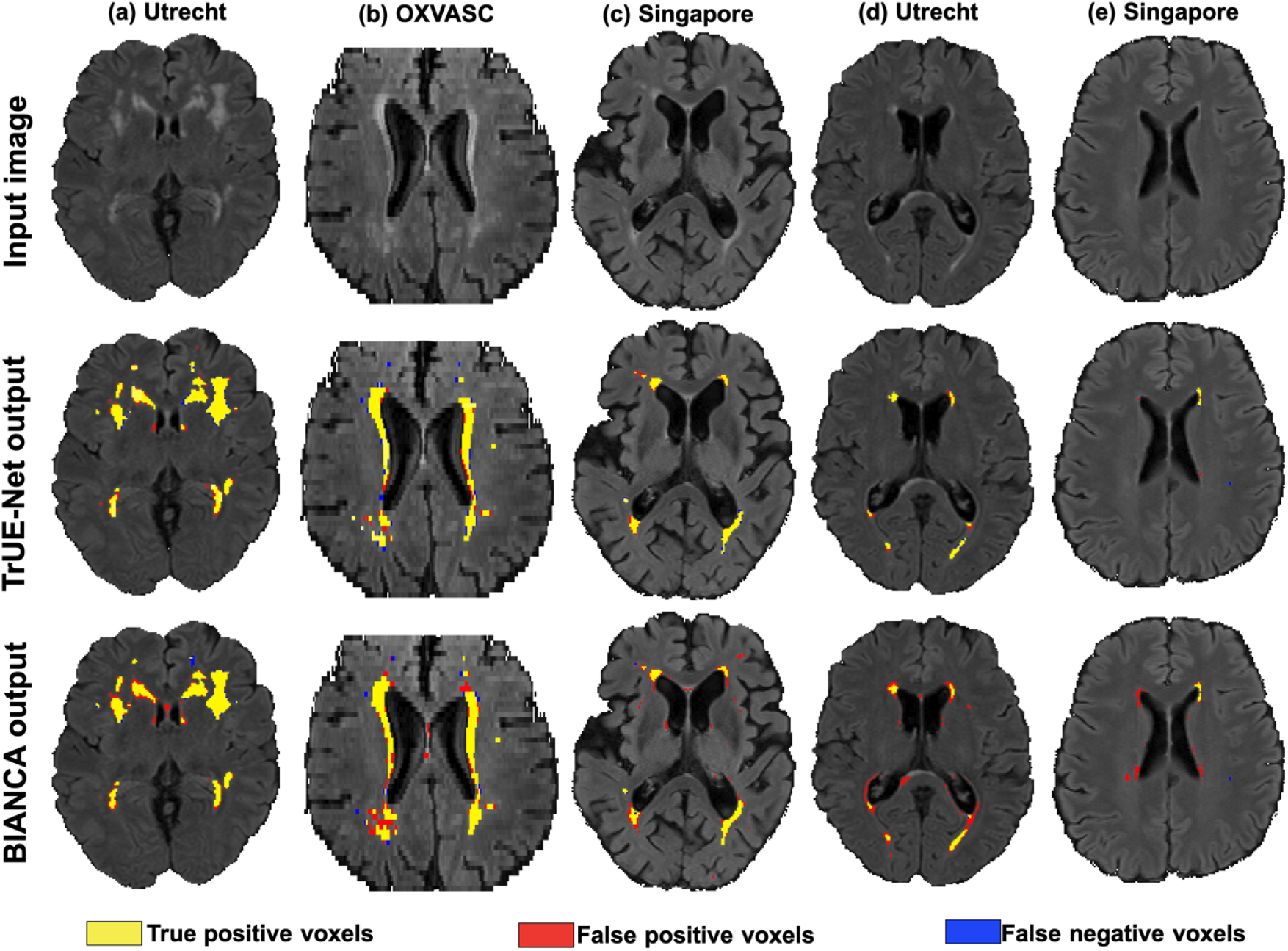
Sample results for comparison of TrUE-Net segmentation with those of BIANCA. Left to right: decreasing order of lesion load from the Utrecht (a,d), Singapore (c, e) and OXVASC (b) datasets. True positive, false positive and false negative voxels are indicated in yellow, red and blue colour respectively.

Overall, TrUE-Net outperforms BIANCA in both voxel-wise and cluster-wise metrics. Figure 11 shows the boxplots comparing the performance metrics between TrUE-Net and BIANCA. The corresponding performance values and p-values of the paired t-tests are reported in table 3. TrUE-Net provides significantly better results than BIANCA for almost all the performance metrics. BIANCA provided the worst performance on the MWSC dataset, with both more false positives and false negatives than TrUE-Net. In the NDGEN dataset, the voxel-wise TPR values are not significantly different for TrUE-Net and BIANCA, indicating that the two methods performs equally on this dataset. Also voxel-wise FPR were not significantly different between the two methods in both NDGEN and OXVASC datasets.

**Figure 11:**
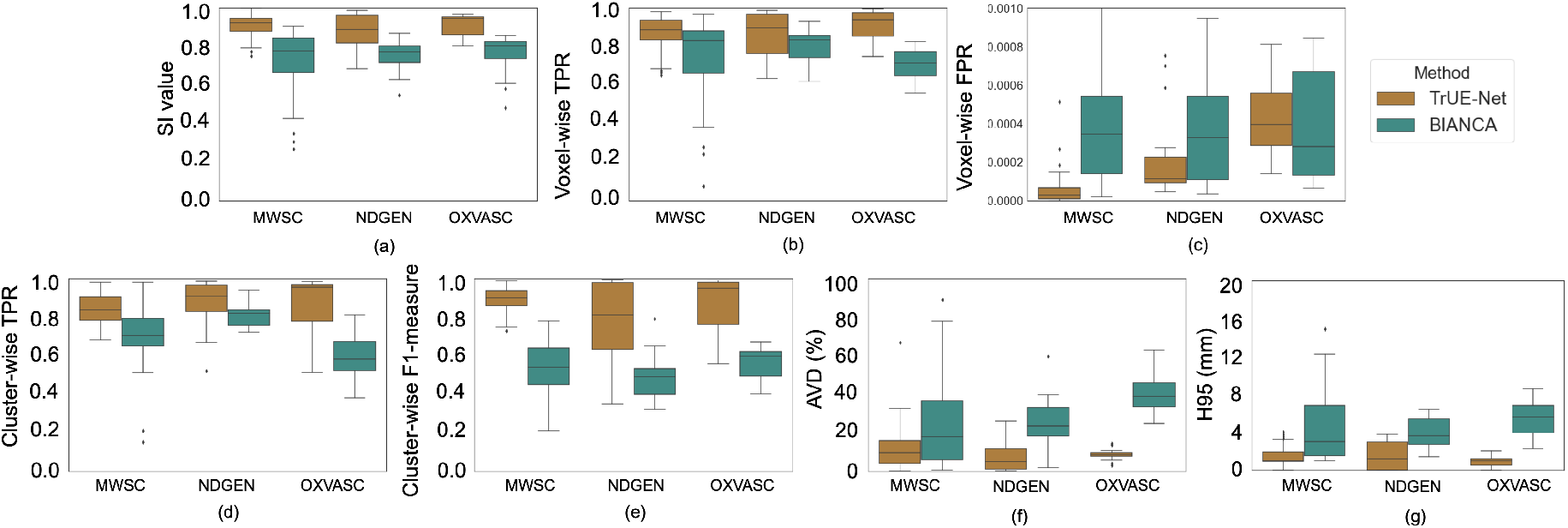
Boxplots of performance metrics obtained for TrUE-Net and BIANCA on the Utrecht, Singapore, Amsterdam, NDGEN and OXVASC datasets - (a) SI value, (b) Absolute volume difference (AVD), (c) voxel-wise TPR, (d) voxel-wise FPR, (e) cluster-wise TPR, (f) cluster-wise F1-measure and (g) 95^th^ percentile of Hausdorff distance.

**Table 3:** Comparison of TrUE-Net performance with BIANCA, along with p-values of two-tailed paired t-test results on the MWSC, NDGEN and OXVASC datasets (the best performance values highlighted in bold).

|  |  | MWSC | NDGEN | OXVASC |
| --- | --- | --- | --- | --- |
| SI | TrUE-Net | $0.91 \pm 0.06$ | $0.88 \pm 0.09$ | $0.91 \pm 0.06$ |
| | BIANCA | $0.73 \pm 0.15$ | $0.76 \pm 0.08$ | $0.75 \pm 0.11$ |
|  | p-value | **$\ll 0.001$** | **$< 0.001$** | **$< 0.001$** |
| Voxel-wise<br>TPR | TrUE-Net | $0.87 \pm 0.09$ | $0.86 \pm 0.12$ | $0.91 \pm 0.08$ |
| | BIANCA | $0.74 \pm 0.20$ | $0.80 \pm 0.08$ | $0.71 \pm 0.08$ |
|  | p-value | **$< 0.001$** | 0.10 | **$< 0.001$** |
| Voxel-wise<br>FPR | TrUE-Net | $0.7 \times 10^{-5}$ | $4.1 \times 10^{-5}$ | $4.2 \times 10^{-5}$ |
| | BIANCA | $4.1 \times 10^{-4}$ | $5.6 \times 10^{-4}$ | $5.9 \times 10^{-4}$ |
|  | p-value | **$\ll 0.001$** | 0.07 | 0.64 |
| Cluster-wise<br>TPR | TrUE-Net | $0.85 \pm 0.08$ | $0.87 \pm 0.12$ | $0.85 \pm 0.18$ |
| | BIANCA | $0.70 \pm 0.14$ | $0.81 \pm 0.06$ | $0.60 \pm 0.12$ |
|  | p-value | **$\ll 0.001$** | 0.06 | **$< 0.001$** |
| Cluster-wise<br>F1-measure | TrUE-Net | $0.89 \pm 0.06$ | $0.78 \pm 0.21$ | $0.87 \pm 0.15$ |
| | BIANCA | $0.54 \pm 0.13$ | $0.47 \pm 0.12$ | $0.57 \pm 0.08$ |
| | p-value | $\approx 0$ | **$< 0.001$** | **$< 0.001$** |
| AVD (%) | TrUE-Net | $11.9 \pm 11.4$ | $8.1 \pm 9.0$ | $8.5 \pm 3.0$ |
| | BIANCA | $36.9 \pm 70.24$ | $25.1 \pm 13.1$ | $40.9 \pm 10.7$ |
|  | p-value | **0.007** | **$< 0.001$** | **$\ll 0.001$** |
| H95 (mm) | TrUE-Net | $1.5 \pm 1.1$ | $1.5 \pm 1.4$ | $0.9 \pm 0.7$ |
| | BIANCA | $4.9 \pm 5.2$ | $3.8 \pm 1.5$ | $6.1 \pm 3.7$ |
|  | p-value | **$< 0.001$** | **$< 0.001$** | **$< 0.001$** |

### 3.5. Comparison with the top ranking method of MWSC 2017

Figure 12 shows the boxplots comparing the performance metrics between TrUE-Net and the method proposed in Li et al. (2018). The corresponding values and p-values of the paired t-tests are reported in table 4. TrUE-Net achieves significantly higher SI values compared to Li et al. (2018). However, Li et al. (2018) achieves better cluster-wise TPR values indicating that the method detects more true-lesions compared to TrUE-Net. On the other hand, the cluster-wise F1-measure value is significantly higher for TrUE-Net, which shows that Li et al. (2018) also detects more false positive clusters, while TrUE-Net provides more cluster-wise precision, resulting in a higher cluster-wise F1-measure value.

**Figure 12:**
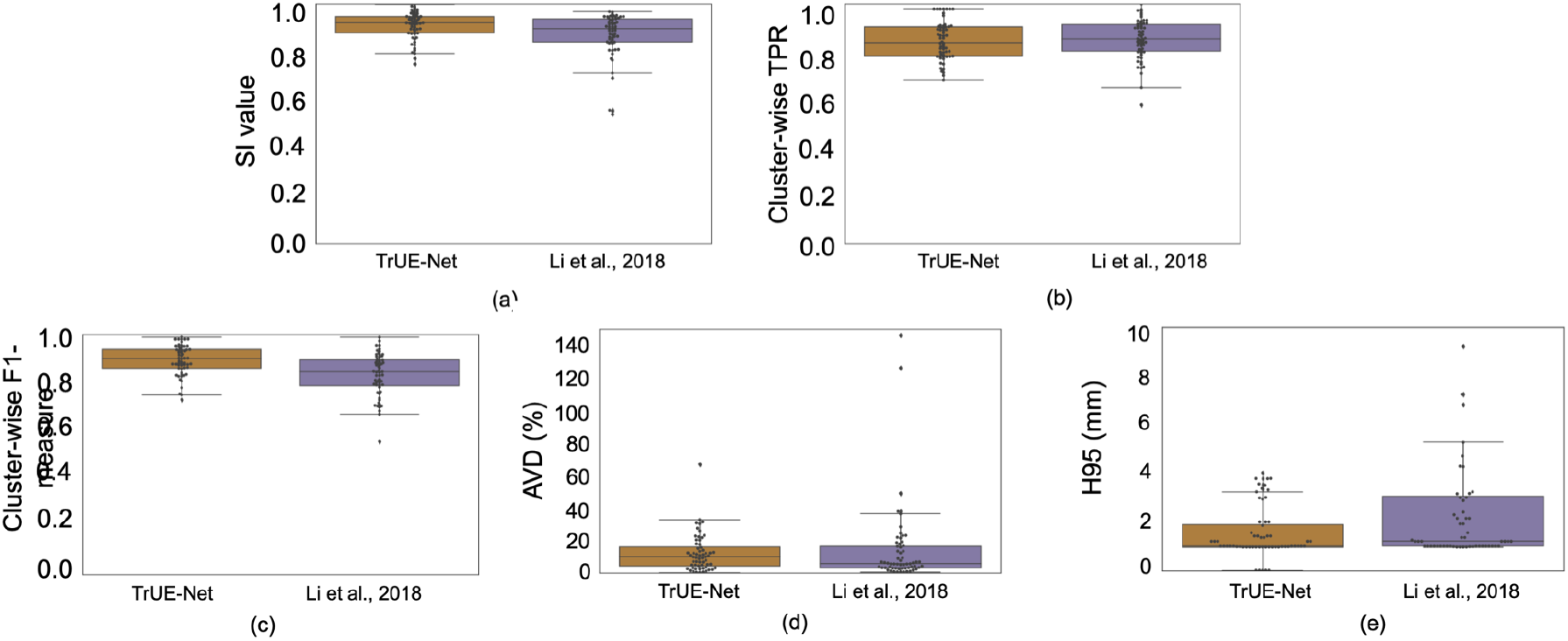
Boxplots of performance metrics obtained for TrUE-Net and Li et al. (2018) on the MWSC dataset - (a) SI value, (b) Cluster-wise TPR, (c) Cluster-wise F1-measure, (d) Absolute volume difference (AVD) and (e) 95^th^ percentile of Hausdorff distance.

**Table 4:** Comparison of LOO results of TrUE-Net with those reported in Li et al. (2018) on the same set of subjects from the MWSC dataset (the best value for each measure is highlighted in bold).

| Perf. metrics | TrUE-Net | Li et al., 2018 | p-value |
| --- | --- | --- | --- |
| SI | **0.91 ± 0.06** | 0.87 ± 0.1 | **0.0045** |
| Cluster-wise TPR | 0.84 ± 0.08 | **0.85 ± 0.08** | 0.70 |
| Cluster-wise F1 measure | **0.89 ± 0.06** | 0.83 ± 0.09 | **< 0.001** |
| AVD (%) | **11.6 ± 11.4** | 13.8 ± 25.17 | 0.58 |
| H95 (mm) | **1.5 ± 1.1** | 3.6 ± 5.93 | **0.008** |

### 3.6. Comparison with other existing methods

In order to contextualise the impact of the methods/improvements presented in this work, table 5 illustrates an indirect comparison with other existing methods in the literature. Among the non-DL methods, the performance is mostly high for supervised methods, with anatomy and appearance-based features used in addition to intensity. The SI values obtained with TrUE-Net are on par with the values reported in previous studies. Also, the performance of TrUE-Net is comparable to the top-performing DL methods in the challenge. However, it is not a direct comparison since the methods from the challenge were evaluated on unseen test data, and also excluding the labels corresponding to other pathologies. As future work we will submit our TrUE-Net model to the challenge in order to achieve a more direct comparison with the other approaches that took part in the challenge. Once the method is submitted to the challenge, a docker image will be available to download from the challenge webpage (http://wmh.isi.uu.nl/).

**Table 5:** Comparison of existing methods (including BIANCA) with TrUE-Net.

| Method | Type<sup>1</sup> | Population - study<sup>2</sup><br>(subjects) | Image modalities | Dice | Sens<sup>3</sup> | Spec<sup>4</sup> |
| --- | --- | --- | --- | --- | --- | --- |
| Wang et al. (2012) | I,U | Singapore ageing cohort (272) | T1, T2, FLAIR | 0.77 | 0.81 | 0.97 |
| Gibson et al. (2010) | I,U | WM disease (18) | FLAIR | 0.81 | - | - |
| De Boer et al. (2009) | I,S | HC - Rotterdam scan study(6) | T1, PD, FLAIR | 0.72 | 0.79 | - |
| Steenwijk et al. (2013) | I,S | Hypertension (20) | T1, FLAIR | 0.84 | - | - |
| Damangir et al. (2012) | IA,S | HC, Alzheimer's, dementia - DemWest (102) | T1, FLAIR | - | 0.90 | 0.99 |
| Ghafoorian et al. (2016) | IA,S | Small vessel disease dataset (50) | PD, T2, FLAIR | - | 0.73 | - |
| Yoo et al. (2014) | I,S | Longitudinal/dementia - Korean study (32) | FLAIR | 0.76 | - | - |
| Jeon et al. (2011) | IA,U | SVD - AMPETIS (45) | PD, T2, FLAIR | 0.90 | - | - |
| Yang et al. (2010) | IA,U | Dementia - LEILA (30) | FLAIR | 0.81 | - | - |
| Shi et al. (2013) | IAAp,U | Acute infarction (91) | DWI, T1, FLAIR | 0.84 | 0.80 | - |
| Samaille et al. (2012) | IAAp,U | MCI, CADASIL (67) | T1, FLAIR | 0.72 | - | - |
| Kruggel et al. (2008) | IAAp | HC, dementia - LEILA (116) |  | - | 0.90 | 0.91 |
| Khademi et al. (2011) | IAAp,U | Subjects with lesions (24) | FLAIR | 0.83 | 0.82 | 0.99 |
| Admiraal-Behloul et al. (2005) | IA,U | Vasc. disease - PROSPER (100) | PD, T2, FLAIR | 0.75 | - | - |
| Anbeek et al. (2004) | IA,S | Arterial vascular disease (20) | T1, IR, PD, T2, FLAIR | 0.80 | 0.97 | 0.97 |
| Li et al. (2018) | DL | MWSC TS | T1, FLAIR | 0.80 | 0.84 | - |
| Andermatt et al. (2016) | DL | MWSC TS | T1, FLAIR | 0.78 | 0.83 | - |
| Berseth | DL | MWSC TS | T1, FLAIR | 0.77 | 0.73 | - |
| Valverde et al. (2017) | DL | MWSC TS | T1, FLAIR | 0.77 | 0.75 | - |
| Kuijf et al. (2019) | DL | MWSC TS | T1, FLAIR | 0.77 | 0.59 | - |
| Kuijf et al. (2019) | DL | MWSC TS | T1, FLAIR | 0.72 | 0.63 | - |
| Xu et al. (2017) | DL | MWSC TS | T1, FLAIR | 0.73 | 0.63 | 0.67 |
| Ghafoorian et al. (2017) | DL | Elder SVD (50) | T1, FLAIR | 0.78 | - | - |
| BIANCA (Griffanti et al., 2016) | IAAp,S | NDGEN (21) | T1, FLAIR | 0.77 | 0.80 | - |
|  |  | OXVASC (18) | T1, FLAIR, MD | 0.74 | 0.71 | - |
|  |  | MWSC TrS (60) | T1, FLAIR | 0.73 | 0.74 | - |
| TrUE-Net | DL | MWSC TrS (60) | T1, FLAIR | 0.91 | 0.87 | - |
|  |  | NDGEN (20) | T1, FLAIR | 0.88 | 0.86 | - |
|  |  | OXVASC (18) | T1, FLAIR | 0.91 | 0.91 | - |

<sup>1</sup> Type: I - intensity, IA - intensity + anatomy, IAAp - intensity + anatomy + appearance, U - unsupervised, S - supervised, DL - deep learning
<sup>2</sup> Population-study: MWSC TS - MICCAI WMH segmentation Challenge test dataset, MWSC TrS - MICCAI WMH segmentation Challenge training dataset (Kuijf et al., 2019)
<sup>3</sup> Sensitivity or Voxel-wise TPR, <sup>4</sup> Specificity.

## 4. Discussion and conclusions

In this work, we proposed a DL model using an ensemble of U-Nets, named TrUE-Net, for accurate WMH segmentation. First, we investigated the effect of various training hyperparameters on model optimisation. We then studied the effect of various components of the loss function on the segmentation performance. On data from 5 cohorts, we evaluated the overall segmentation performance as well as the performance in deep and periventricular regions separately. In addition, we directly compared our method with a non-DL method (BIANCA) and a DL method (the top ranking method of MWSC 2017). Finally, we provided an the indirect comparison with various methods proposed in the literature.

When optimising our model, we observed that using a lower batch size resulted in noisy but lower loss values. The lower batch size could be advantageous due to two reasons: lower computation load per batch (hence higher speed) and faster convergence due to the regularisation effect of noisy gradient estimation (Wilson and Martinez, 2003; Keskar et al., 2016). Firstly, using smaller batches reduces the gradient estimation time per iteration. However, this would increase the overall number of iterations per epoch, which brings us to the second advantage. Due to the noisy estimate of mean gradient over batches, during subsequent iterations in our experiments we observed that the cost function fluctuates, getting out of some spurious local minima and converging to a better local minima quickly when using smaller batches. Additionally, in our experiments smaller batch sizes avoided over-fitting, as evident from the lower validation loss for a batch size of 8, compared to 16 and 32 (figure 3a). Regarding the parameter *ϵ*, we chose the value 1 × 10^−4^ as an optimal value for further experiments, since it showed lower loss values. The parameter *ϵ* is used in the denominator (to avoid divide-by-zero error) in the determination of updates for weights. Having a very low *ϵ* value results in estimation of larger weight updates, leading to unstable optimisation as shown in the case of 1 × 10^−6^ (figure 3b). In the case of learning rate, choosing a higher value results in earlier convergence with higher loss values. On the other hand, lower values require more epochs to converge due to smaller steps of the updates in weights. Hence, we chose an optimal learning rate schedule between 1 × 10^−3^ and 1 × 10^−6^, figure 3c, for our further experiments.

When comparing the results using different components of the loss function, we observed that the CE loss component is responsible for identification of the majority of lesions against the background voxels, while the Dice loss component is more sensitive to smaller lesions and precise boundaries. We observed that weighting the CE loss with values based on distances from the ventricles and GM imparts more anatomical information, and additionally addresses the class imbalance. This avoids over-segmentation of highly bright PWMHs and under-segmentation of DWMHs (figure 5c,d). In addition, using the Dice loss component also results in the detection of more subtle deep lesions, giving more accurate segmentation (figure 5e) and avoiding over-segmentation in low lesion load subjects. This results in higher SI values in these low lesion load subjects (shown in figure 6).

The TrUE-Net model gave good performance in all 5 datasets. In particular, in the MWSC dataset, TrUE-Net achieved the best cluster-wise F1-measure and the lowest voxel-wise FPR indicating precise and accurate segmentation. The MWSC cohort consists of subject with variations in lesion load and acquisition characteristics. Even though the SI value on the NDGEN dataset is lower than the other two datasets, TrUE-Net detects more small true lesions (best cluster-wise TPR) compared to the other datasets.

When comparing the performance of our model in the deep and the periventricular regions, the trend of performance metrics for DWMHs and PWMHs remained consistent across datasets. In general, we observed that SI values and voxel-wise TPR values were higher for PWMHs when compared to DWMHs. In the deep regions, cluster-wise TPR was higher than that in the periventricular regions. This means that more DWMHs are detected correctly (also evident from higher cluster-wise F1-measure values). However, the lesion boundaries are delineated better in PWMHs when compared to DWMHs, as indicated by slightly higher Hausdorff distance for DWMHs (figure 9 and table 2). At this point, it is worth remembering that manual segmentation is our gold standard but not necessarily the absolute truth. Therefore these errors in the delineation of DWMHs could be due to inconsistencies in manual segmentation (due to low contrast and other confounders) rather than lower model sensitivity.

On comparing TrUE-Net with BIANCA, we found that TrUE-Net outperforms BIANCA with significant differences in the performance metrics in almost all datasets. Overall, we found that TrUE-Net achieves the highest SI values, cluster-wise F1-measures and the lowest Hausdorff distance values for all datasets, and provides better segmentation in various lesion load cases (figure 11). TrUE-Net performs well even in subjects with low lesion load, detecting fewer false positives than BIANCA. Also, TrUE-Net provides results comparable with the top ranking method of the MICCAI Challenge, with significantly higher mean SI value and cluster-wise F1-measure. When performing an indirect comparison of TrUE-Net with other existing methods, we observed that only a few methods have been evaluated on a variety of datasets (pathological population and/or healthy subjects). Most of the methods considered for the comparison analysis are tested on datasets from a specific population, and hence might require additional experimentation to validate/improve their generalisability (e.g. fine-tuning of parameters, change of cut-off/threshold values). Some of the multimodal methods in the literature allow the flexibility of choosing different input modalities, while others use fixed sets of modalities. There are both pros and cons associated with using either of the methods. Methods that are flexible allow users to use various available modalities and could handle datasets with missing modalities. On the other hand, the choice of input modalities becomes an additional parameter to tune when applied on an unknown dataset. Also, from our comparison analysis we observed that the use of more modalities might not always lead to better performance. For instance, for the OXVASC dataset, the best performance for BIANCA was achieved with T1 + FLAIR + MD, while TrUE-Net provided better SI values by using only T1 + FLAIR.

In conclusion, we proposed a model that provides accurate segmentation, with better performance than BIANCA and on par with the top ranking method of MWSC 2017. We evaluated it on various datasets with different population and lesions characteristics. Regarding the tool availability, TrUE-Net is currently available in its python implementation in https://git.fmrib.ox.ac.uk/vaanathi/true_net_wmh_segmentation_pytorch and will be integrated as an independent WMH segmentation tool in a future release of FSL.

## 5. Acknowledgements

This work was supported by the Engineering and Physical Sciences Research Council (EPSRC), Medical Research Council (MRC) [grant number EP/L016052/1] and Wellcome Centre for Integrative Neuroimaging, which has core funding from the Wellcome Trust (203139/Z/16/Z). The computational aspects of this research were funded from National Institute for Health Research (NIHR) Oxford BRC with additional support from the Wellcome Trust Core Award Grant Number 203141/Z/16/Z. The Oxford Vascular Study is funded by the National Institute for Health Research (NIHR) Oxford Biomedical Research Centre (BRC), Wellcome Trust, Wolfson Foundation, the British Heart Foundation and the European Union’s Horizon 2020 programme (grant 666881, SVDs@target). VS is supported by the Wellcome Centre for Integrative Neuroimaging. GZ is supported by the Italian Ministry of Education (MIUR) and by a grant “Dipartimenti di eccellenza 2018-2022”, MIUR, Italy, to the Department of Biomedical, Metabolic and Neural Sciences, University of Modena and Reggio Emilia. PMR is in receipt of a NIHR Senior Investigator award. The views expressed are those of the author(s) and not necessarily those of the NHS, the NIHR or the Department of Health. MJ is supported by the NIHR Oxford Biomedical Research Centre (BRC). LG is supported by the Oxford Parkinson’s Disease Centre (Parkinson’s UK Monument Discovery Award, J-1403), the MRC Dementias Platform UK (MR/L023784/2), and the National Institute for Health Research (NIHR) Oxford Health Biomedical Research Centre (BRC).

We acknowledge all the participants. For the NDGEN dataset, we are grateful to Prof. Gordon K. Wilcock and all the staff of Oxford Project to Investigate Memory and Ageing (OPTIMA) study. For the OXVASC dataset, we acknowledge the use of the facilities of the Acute Vascular Imaging Centre, Oxford. We also thank Dr. Chiara Vincenzi and Dr. Francesco Carletti for their help on generating the manual masks used in our experiments.

MJ and LG receive royalties from licensing of FSL to non-academic, commercial parties. The authors report no potential con icts of interest.

